# Metagenomic Next-Generation Sequencing of Rectal Swabs for the Surveillance of Antimicrobial Resistant Organisms on the Illumina Miseq and Oxford MinION Platforms

**DOI:** 10.1101/2020.04.16.044214

**Authors:** Rebecca Yee, Florian P. Breitwieser, Stephanie Hao, Belita N.A. Opene, Rachael E. Workman, Pranita D. Tamma, Jennifer Dien-Bard, Winston Timp, Patricia J. Simner

**Affiliations:** Department of Pathology, Johns Hopkins University School of Medicine, Baltimore, MD; Center for Computational Biology, McKusick-Nathans Institute of Genetic Medicine, Johns Hopkins School of Medicine, Baltimore, MD; Department of Biomedical Engineering, Johns Hopkins University, Baltimore, MD; Department of Pediatrics, Johns Hopkins University School of Medicine, Baltimore, MD; Department of Pathology and Laboratory Medicine, Children’s Hospital of Los Angeles and Keck School of Medicine at the University of Southern California, Los Angeles, California, USA

**Keywords:** metagenomics, next-generation sequencing, antimicrobial resistance, microbiome, resistome, surveillance

## Abstract

**Purpose:** Antimicrobial resistance (AMR) is a public health threat where efficient surveillance is needed to prevent outbreaks. Existing methods for detection of gastrointestinal colonization of multidrug-resistant organisms (MDRO) are limited to specific organisms or resistance mechanisms. Metagenomic next-generation sequencing (mNGS) is a more rapid and agnostic diagnostic approach for microbiome and resistome investigations. We determined if mNGS can detect MDRO from rectal swabs in concordance with standard microbiology results.

**Methods:** We performed and compared mNGS performance on short-read Illumina MiSeq (N=10) and long-read Nanopore MinION (N=4) platforms directly from peri-rectal swabs to detect vancomycin-resistant enterococci (VRE) and carbapenem-resistant Gram-negative organisms (CRO).

**Results:** We detected *E. faecium* (N=8) and *E. faecalis* (N=2) with associated *van* genes (9/10) in concordance with VRE culture-based results. We studied the microbiome and identified CRO organisms, *P. aeruginosa* (N=1), *E. cloacae* (N=1), and KPC-producing *K. pneumoniae* (N=1). Nanopore real-time detection detected the *bla_KPC_* gene in 2.5 minutes and provided genetic context (*bla*_KPC_ harbored on pKPC_Kp46 IncF plasmid). Illumina sequencing provided accurate allelic variant determination (i.e., *bla*_KPC-2_) and strain typing of the *K. pneumoniae* (ST-15). Conclusions: We demonstrated an agnostic approach for surveillance of MDRO, examining advantages of both short and long-read mNGS methods for AMR detection.

## INTRODUCTION

Early detection of colonization by multidrug-resistant organism (MDRO) such as vancomycin-resistant enterococci (VRE) and carbapenem-resistant organisms (CRO) can lead to early implementation of infection prevention practices, antimicrobial optimization, and prevention of invasive infections [1]. Current methods (selective culture techniques or PCR) are targeted towards a specific MDRO [2]. To identify novel mechanisms of resistance or emerging pathogens, a broader approach to detect and characterize MDRO is required.

Metagenomic next-generation sequencing (mNGS) of specimens using next-generation sequencing (NGS) platforms is an agnostic approach. mNGS amplifies any DNA in the sample. Thus, this approach can query the entire microbiome of the sample, and provide valuable information about the resistome (all known antimicrobial resistance genes), and plasmids [3].

In this proof-of-concept study, we determined if mNGS using short-read Illumina sequencing and long-read Oxford Nanopore sequencing can be applied to rectal swabs for detection of VRE and CROs identified by standard microbiology results.

## Materials and Methods

### Bacterial Isolates and Characterization

Remnant rectal surveillance swabs from 10 ICU patients hospitalized at Johns Hopkins Hospital were evaluated for VRE on VRE Select chromogenic agar and CRO by the Direct MacConkey method [2]. Carbapenemase production was detected by the Carba NP assay and Check-MDR CT103XL from cultured isolates and Check-Direct CPE screen assay for the BD MAX instrument from rectal swabs (Check-Points; Becton Dickinson) [4]. Bacteria were identified by matrix-assisted laser-desorption ionization time-of-flight mass spectrometry (Bruker Daltonic). Positive controls (10^2^, 10^4^, or 10^6^ CFU/mL) swabs seeded with *Klebsiella pneumoniae* ATCC BAA-1705 (*bla_KP_C*), *Enterococcus faecalis* ATCC 51299 (*vanB*) and *Enterococcus faecium* ATCC 700228 (*vanA*) and negative control swabs were also sequenced. ESwabs were de-identified, frozen at −70°C, and DNA was extracted from the broth (500 μl) using the Zymo ZR Fungal/Bacterial Miniprep Kit.

### Illumina Sequencing

Library preparation was performed using Illumina NexteraXT^™^ DNA Sample Prep Kit per manufacturer’s protocol followed by AMPure XP (Beckman Coulter) purification. Normalized samples were pooled (n=4) and sequenced on a MiSeq v3 2×75 flowcell. Reads were assembled using metaSPAdes [5].

### Oxford Nanopore Sequencing

Library preparation was performed using the low-input genomic DNA sequencing kit protocol for SQK-MAP006 per manufacturer’s instructions. Libraries were loaded onto a R7.3 flowcell, sequenced using MinKNOW, and basecalled using Metrichor. Only Nanopore 2d high quality reads were used.

### Bioinformatics

Taxonomic analysis was performed with Kraken and plotted using Krona [6]. Resistance genes were queried using BLAST and against ResFinder using Abricate [5]. Alignment to *K. pneumoniae* strain KPNIH49 (RefSeq GCF_002903025.1) was done using bowtie2 and visualized using Pavian [7]. Data analysis and heatmap visualization was done using R statistical environment [8]. Our sequencing data, scrubbed of human reads, has been deposited at SRA (accession pending).

## Results

### Characterization of Microbiome from Rectal Swabs

The results from both platforms were concordant with standard microbiology culture methods (Table 1). We detected VRE (*E. faecium* and/or *E. faecalis*)*, vanA* and/or *vanB* in all samples, and predominant CRO organisms (*E. cloacae, P. aeruginosa and K. pneumoniae*) in 3 samples (Figure 1A-B). Both platforms detected the same or similar species as the top three dominant organisms (Table 1).

**Fig. 1.**
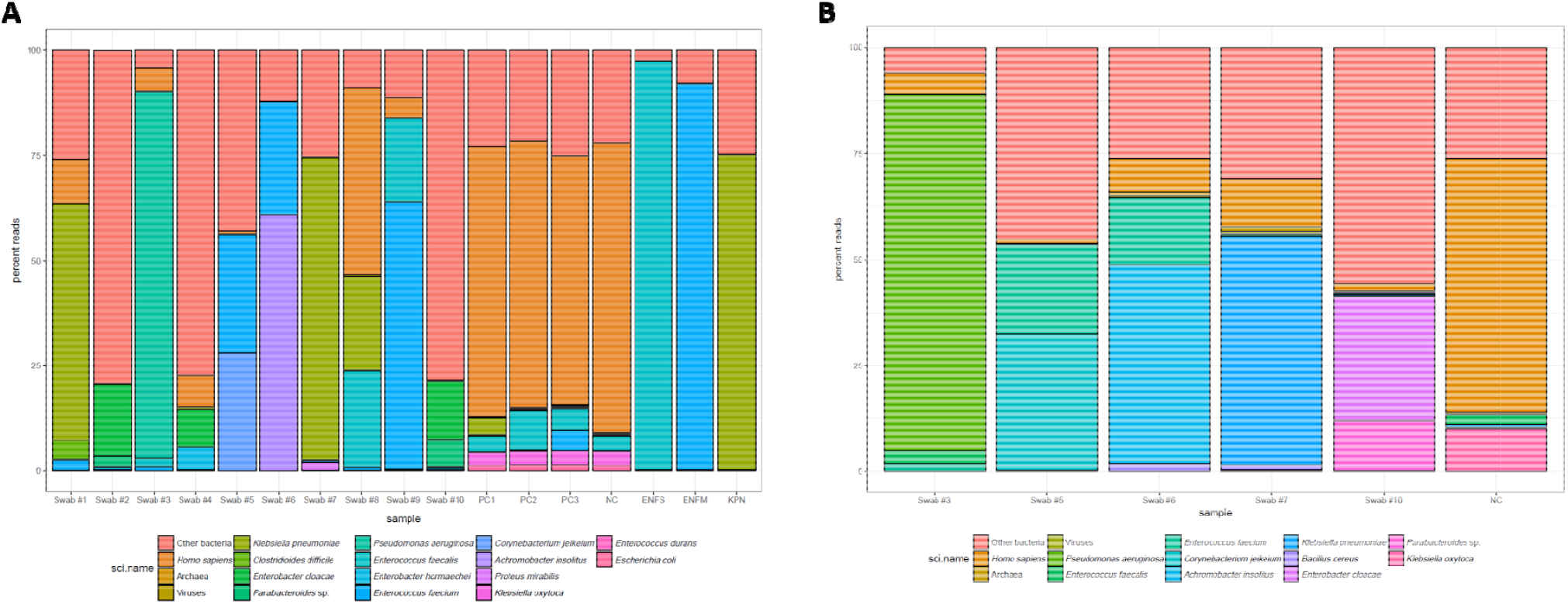
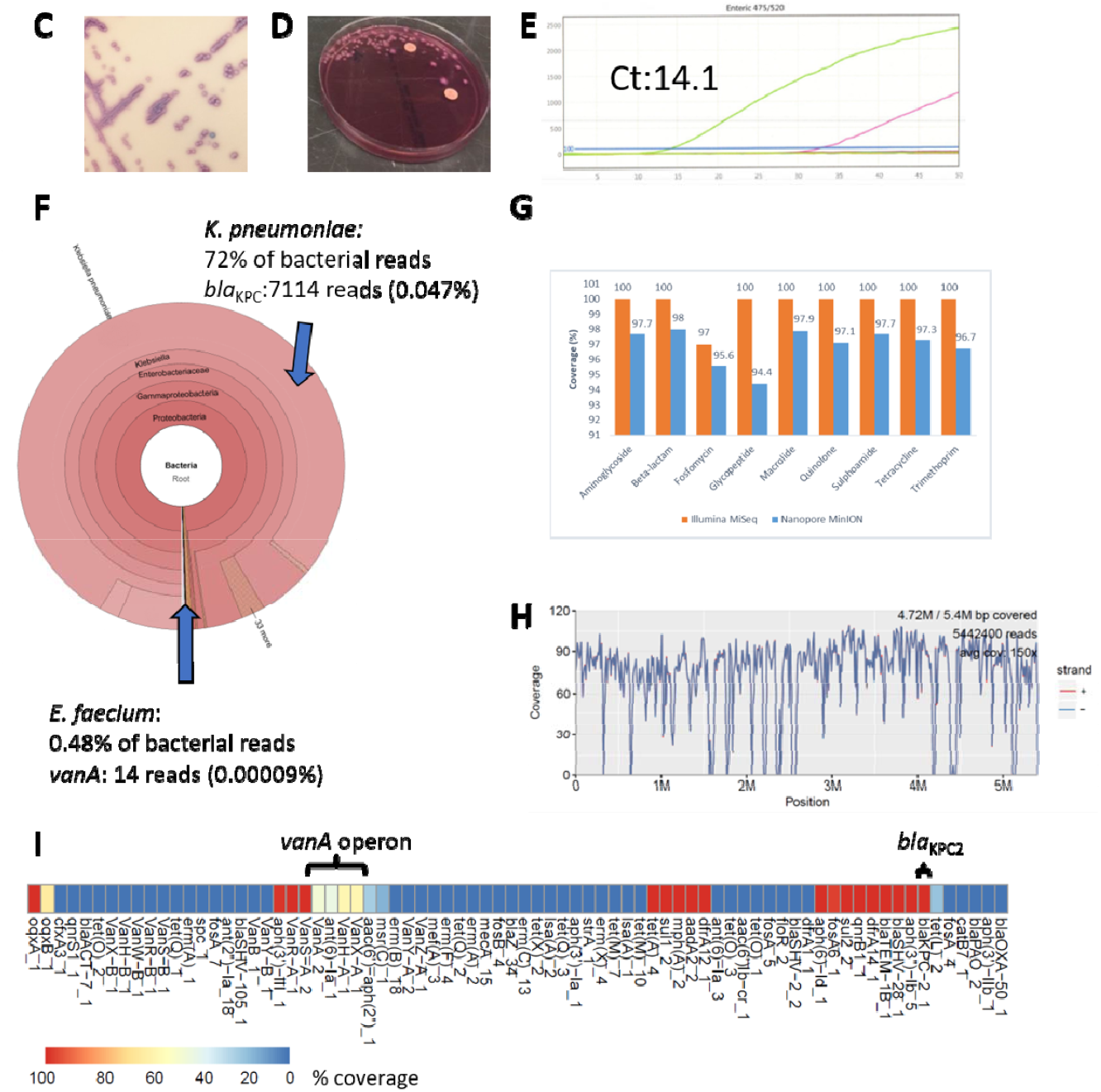
Microbiome and Resistome Analyses Performed on Rectal Swabs. Depiction of organisms with the most percentage of reads from all ten swabs tested on (A) Illumina MiSeq and selected swabs tested on (B) Oxford Nanopore MinION. Taxonomic analysis was performed with Kraken using kmer sizes of 24. Positive controls (PC1, PC2, PC3) were spiked with varying proportions (10^2^, 10^4^ or 10^6^ CFU/mL) of the following organisms *Klebsiella pneumoniae (bla_KPC_* positive; KPN), *Enterococcus faecalis (vanB* positive; ENFS) and *Enterococcus faecium (vanA* positive; ENFM). Organisms used for the positive control spike-ins were also sequenced individually. Negative control (NC) was a pool of rectal E-swabs found to be negative for VRE and CRO by culture. Culture-based methods showed rectal swab #7 contained a (C) positive vancomycin-resistance *Enterococcus faecium* on VRE Select chromogenic agar and (D) 1+ growth of a lactose fermenter producing a 12 mm zone of inhibition around an ertapenem disk on MacConkey agar. (E) Carbapenemase gene *bla*_KPC_ was detected with a Ct value of 14.1 using CheckDirect CPE screen assay directly from the rectal swab. (F) Krona plot of metagenomics analyses from sequencing performed on Illumina MiSeq revealed *K. pneumoniae* as the dominant organism and also detection of *E. faecium* as the second organism, both of which were detected by culture-based methods. (G) Coverage comparison of short-read Illumina MiSeq and long-read Oxford Nanopore MinION revealed higher accuracy and coverage on Illumina MiSeq. (H) Accuracy of Illumina reads allowed for straining typing of the KPC-producing *K. pneumoniaeas* ST-15. (I) Resistome analyses demonstrated the presence of both vancomycin (*vanA* operon) and carbapenem resistance (*bla*_KPC_) genes based on adaptation of using ResFinder and additional BLAST analyses

**Table 1:** Summary of Culture Based Results for the Detection of Antimicrobial Resistant Organisms from Rectal Swabs Compared to mNGS on the MiSeq and MinION Platforms

| Study # | Vancomycin-resistant enterococci (VRE) Culture Results |  | Carbapenem-Resistant Organism Culture Results |  | CheckDirect CPE Results | Miseq Data |  | Nanopore Data |  |
| --- | --- | --- | --- | --- | --- | --- | --- | --- | --- |
|  | VRE culture results | Organism based on chromogen | Gram-negative growth on MacConkey Agar with ertapenem disks<sup>a</sup> | Carbapenem resistant organism (CRO) culture results | Detection of carbapenemase genes directly from rectal swabs | Dominant Bacterial organisms | Detection of *vanA*, *vanB* or *bla*<sub>KPC</sub> | Dominant Bacterial Organisms<sup>b</sup> | Detection of *vanA*, *vanB* or *bla*<sub>KPC</sub><sup>b</sup> |
| 1 | Positive for VRE | *E. faecium* | 4 + Lactose fermenter; >27mm zone diameter | Negative | Negative | *Klebsiella pneumoniae*,<br>*Clostridioides difficile*,<br>*Enterococcus faecium* | *vanA* positive | N/A | N/A |
| 2 | Positive for VRE | *Enterococcus faecalis* | 4+ Mixed lactose fermenters; >27mm zone diameter | Negative | Negative | *Enterobacter cloacae*,<br>*Parabacteroides distasonis*,<br>*Enterobacter asburiae* | *vanA* and *vanB* positive | N/A | N/A |
| 3 | Positive for VRE | *E. faecium* | 4+ Non-lactose fermenter; no zone around ertapenem disk | Positive: Meropenem resistant<br>*Pseudomonas aeruginosa* | Negative | *Pseudomonas aeruginosa*,<br>*Enterococcus faecalis*,<br>*Enterococcus faecium* | *vanA* positive | *Pseudomonas aeruginosa*,<br>*Enterococcus faecalis*,<br>*Enterococcus faecium* | *vanA* positive |
| 4 | Positive for VRE | *E. faecium* | 4+ Non-lactose fermenter; 21 mm zone around ertapenem disk | Positive: Ertapenem resistant<br>*Enterobacter cloacae* | Negative | *Enterobacter cloacae*,<br>*Enterobacter asburiae*,<br>*Cronobacter sakazakii* | no detection | N/A | N/A |
| 5 | Positive for VRE | *E. faecium* | No Growth | Negative | Negative | *Enterococcus faecium*,<br>*Corynebacterium jeikeium*,<br>*Parabacteroides* spp. | *vanA* positive | *Enterococcus faecium*,<br>*Corynebacterium jeikeium*,<br>*Parabacteroides* spp. | *vanA* positive |
| 6 | Positive for VRE | *E. faecium* | 1+ Non-lactose fermenter; >40 mm zone around ertapenem disk | Negative | Negative | *Achromobacter xylosoxidans*,<br>*Enterococcus faecium*,<br>*Pseudomonas aeruginosa*, | *vanA* positive | *Achromobacter xylosoxidans*,<br>*Enterococcus faecium*,<br>*Bacillus cereus* | *vanA* positive |
| 7 | Positive for VRE | *E. faecium* | 1+ Lactose fermenter; 12 mm zone around ertapenem disk | Positive: KPC-producing *K. pneumoniae* | *bla*<sub>KPC</sub><br>Ct: 14.1 | *Klebsiella pneumoniae*,<br>*Proteus mirabilis*,<br>*Enterococcus faecium* | *vanA*, *bla*<sub>KPC</sub> positive | *Klebsiella pneumoniae*,<br>*Enterococcus faecium*, *Bacillus cereus* | *vanA*, *bla*<sub>KPC</sub> positive |
| 8 | Positive for VRE | *E. faecium* | Mixed Lactose fermenters; 36 mm zone around ertapenem disk | Negative | Negative | *Klebsiella pneumoniae*,<br>*Enterococcus faecalis*,<br>*Enterococcus faecium*, | *vanA* positive | N/A | N/A |
| 9 | Positive for VRE | *E. faecium* | No Growth | Negative | Negative | *Enterococcus faecium*,<br>*Enterococcus faecalis*,<br>*Corynebacterium jeikeium*, | *vanA* positive | N/A | N/A |
| 10 | Positive for VRE | *E. faecalis* | Non-lactose fermenter ("Pseudo"); 39 mm zone around ertapenem | Negative | Negative | *Parabacteroides distasonis*,<br>*Enterobacter cloacae*,<br>*Enterobacter hormaechei* | *vanA*, *vanB*,<br>*bla*<sub>KPC</sub> positive<sup>c</sup> | *Parabacteroides distasonis*,<br>*Enterobacter cloacae*,<br>*Klebsiella pneumoniae* | *vanB* positive |

<sup>a</sup> 1-4+ indicates the amount of growth on the plates. 1+: growth in first quadrant, 2+: is growth in the second quadrant, 3+ is growth
into the third quadrant and 4+ is growth into the 4<sup>th</sup> quadrant.
<sup>b</sup> N/A: Not applicable as swabs 1, 2, 4, 8 and 9 were not run on the Nanopore platform
<sup>c</sup> While *bla*<sub>KPC</sub> was detected in Swab #10, we did not detect *bla*<sub>KPC</sub> using traditional microbiologic methods. Detection was likely due
to errors in de-multiplexing, as Swabs #7 and #10 were ran on the same flowcell. The low number of reads (n=3) for the *bla*<sub>KPC</sub>
gene in Swab #10 suggests that the detection was likely an artifact.

Metagenomic data and frequency of classified reads were similar between both platforms (Figure 1, see web-only Supplementary Table S1). Similar percentages of human reads (range 0.01% and 10%), with Swab #8 as an outlier (44%) were generated. Our negative control had greater than 60% human reads on both platforms.

### AMR Gene Detection

MiSeq detected *vanA* in all samples except Swab #4, which showed a lower quantity of growth by culture suggesting that initial sampling may be low. Swabs #2 and #10 had *vanB* detected and Swab #7 had *bla*_KPC_ detected, corresponding to culture and PCR results from the rectal swab. The *bla*KPC gene was detected by both platforms but non-carbapenemase mediated mechanisms of carbapenem resistance were not confirmed by mNGS. Our bioinformatics analyses also revealed AMR genes for other antimicrobial classes (Figure 1). All controls sequenced generated data concordant with the seeded organisms and resistance genes.

### Case-study: Correlation of Phenotypic Culture Results to Molecular mNGS Analysis

In Swab #7, we identified vancomycin-resistant *E. faecium* (Figure 1C) and a lactose fermenter with a 12 mm zone of inhibition around an ertapenem disk (Figure 1D), further revealed to be a KPC-producing *K. pneumoniae* by PCR from cultured isolate (Figure 1E). From both platforms, we detected *K. pneumoniae* as the dominant and *E. faecium* as the second organism (Figure 1F). MiSeq further determined the allele as *bla*_KPC-2_, identifying mutations to the single nucleotide level for allelic variant identification and strain typing of ST-15 for the predominant *K. pneumoniae* directly from specimen (Figure 1G-H). For other AMR genes, the accuracy and percent coverage for allelic variants detected by MinION (96.9%) was lower than MiSeq (99.7%) (Figure 1G). Meanwhile, MinION’s ability for rapid real-time analysis of AMR genes detected *bla*_KPC_ in 2.3 minutes. Additionally, MinION sequencing provided the ability to associate the *bla*_KPC_ gene to the IncFII-type plasmid pKPC_Kp46 [9]. Heatmap of the full resistome was also generated (Figure 1I).

## Discussion

The shortcomings of targeted PCRs and the requirement for a priori knowledge of the resistance markers is more evident as additional MDROs are identified, highlighting the potential benefits of mNGS as an alternative approach. Despite its high accuracy (99%) in strain typing and AMR allelic variant determination, Illumina sequencing may not be actionable for clinical care usages due to long run times of 24-48 hours. Nanopore sequencing offers advantages of real-time base-calling, detection of resistance genes in as little as 2 minutes and determination of the genetic context of the AMR genes detected. A real-time WGS approach detected all AMR genes within 14 minutes and can shorten time to effective therapy for carbapenem-resistant *K. pneumoniae* infections by 20 h compared to standard approaches [5]. In 6 hours, full annotation of plasmid-based resistance genes was achieved in extended-spectrum β-lactamase-producing *E. coli* and *K. pneumoniae* isolates [10].

Our study here is one of the first to compare platforms for mNGS on rectal swabs where accurate MDRO and resistance genes were identified compared to culture-based methods. Mu *et. al* identified a KPC-producing *K. pneumoniae* isolate using MiSeq but only tested one rectal swab [11]. The similar performance of both platforms seen in our study has been observed in others; comparable phylogenetic trees for *N. gonorrhoeae* [12] and detection of Dengue and Chikungunya viruses from plasma and serum samples [13] have been shown. Similarly, Nanopore sequencing distinguished allelic variants poorly by flagging multiple alleles (i.e., *bla*_KPC_-_2_, *bla*_KPC-3_, etc.) while Illumina sequencing detected single variants (ie., *bla*_KPC-2_) and performed strain typing directly from specimens [14].

Limitations include the lack of broader species and resistance mechanisms tested. Newer sequencing methods (rapid extraction and library kits, Flongle) and flow cells (MinION R.9 and R.10) from Oxford Nanopore may improve accuracy significantly [15].

In conclusion, mNGS analysis may be a promising approach for detection of MDRO from rectal swabs. mNGS allows the study of the entire microbiome, providing important clinical information such as the resistome, allelic variants, and strain typing to guide infection control and patient management. Future studies evaluating newer technologies and automated processes are necessary to increase the efficiency and advance mNGS methods for patient care.

## Notes

### Competing Interest Statement

The authors have declared no competing interest.

